# Temperature and polarity based evolutionary model of the ribosomal complex in *Thermus thermophilus* and *Escherichia coli*

**DOI:** 10.1101/657692

**Authors:** Rashmi Tripathi

**Affiliations:** ConLifeSci, Global Awakening Ideas & Actors (G.A.I.A), Planet number 3, Solar System, Milky Way Galaxy.

## Abstract

The ribosome is considered a molecular fossil of the RNA world and is the oldest molecular machinery of living cells responsible for translating genetic information encoded by messenger RNA(mRNA) to proteins. Currently not much is known regarding how these proteins were assembled and the potential biogeochemical environment that could have shaped their evolution. In order to answer these questions, a comprehensive analysis of the amino acid frequencies of 30S and 50S ribosomal sub-units occurring in thermophile *Thermus thermophilus* and mesophile *Escherichia coli* was performed. The amino acid frequencies in proteins are believed to have been shaped by their pre-biotic abundances in the universe and by heavy bombardment of meteorites on planet earth (4.5-3.8 Ga). Absence of amino acid residues such as cysteine and tryptophan in *T.thermophilus* and *E.coli* proteins hints towards the evolution of small and large subunits prior to the origin of metabolic pathways of amino acid synthesis possibly under anoxic and sulphur free conditions. Moreover, an underrepresentation of readily oxidizable amino acids such as methionine, tyrosine and histidine, indicates that these proteins could have evolved in a more reducing environment as was prevalent on early earth. A comparison of amino acid biases with universal UNIPROT estimates, indicates arginine and lysine overrepresentation, linking a role of these amino acids in ribosomal RNA binding and stabilization corresponding to the RNA world hypothesis whereby RNA molecules drove the assembly of living systems. The continuing prevalence of these amino acid biases in modern proteins reflects the functional stability of ancient proteins constructed during billions of years of evolution and provides glimpses into the evolution of the ancient amino acid code. Step-wise accretion models involving increasing complexity of the amino acid code and the ribosomal sub-units are proposed for *T.thermophilus* and *E.coli*, providing potential insights regarding the origin of ribosomes in a temperature dependent and polar environment.

## Introduction

According to NASA, life is defined as a ‘self-sustaining system capable of Darwinian evolution’. A systems biological perspective that takes into account co-evolving chemistry of the environment and defines ‘life’ as a ‘closed system’ capable of energy capture, growth, reproduction and subsequent evolution seems like a more plausible definition of such complex phenomena. In attempting to model the earliest living forms often referred to as the last universal common ancestors (LUCA)^1,2^, from which arose all major taxa of life including archaea, bacteria and eukaryotes, various attempts have been made to infer the minimal set of protein, ribonucleic acid (RNA) and deoxyribonucleic acid (DNA) molecules that could have constituted the first primitive cells that arose between 3.5 to 0.5 Ga.

Although underlying chemical reactions that could have resulted in the complex biochemistries of living cells remain unknown, speculative evidence regarding the nature of the ‘primordial soup’ has been provided by Miller’s pioneering experiments that simulated volcanic eruption of gases and spark discharge synthesis by lightening^3^. These subsequently led to the identification of ~25 amino acids in the extracts analysed providing a plausible explanation for the evolution of the most common building blocks of life- from simple volatile gases to inorganic minerals serving as potential catalysts^4^. Other proposed sources of synthesis include formation in hydrothermal vents, delivery to earth by icy comets/meteorites via solid phase synthesis and gas phase synthesis in hot molecular cores^5^. Raggi *et al.* propose that more than 80 different amino acids are chemically plausible raising the question regarding the processes that led to selection of ~20 amino acids encoded by the genetic code^6^. One can speculate that the chemical composition of reaction mixtures prevalent in diverse environments could have potentially influenced the chemistry of amino acid and nucleotide containing polymers and that the original last universal common ancestors were more likely a heterogenous population defined by their unique bio-geochemical environments.

Synthesis of complex polymers such as RNAs, DNAs and proteins could have initially utilized energy derived from diverse sources such as spark discharge from lightening, heat, radiation received from the sun and electron transfer reactions employing transition metals such as Fe, Ca and Mg. It is hypothesized that some form of biomineralization process triggered nucleation and growth of these complex bio-polymers that subsequently acquired the ability to replicate and translate information as defined by the molecular dogma of life, i.e DNA <-> RNA->proteins. Thus, arose a complex polymer based system driven by slow reaction kinetics of replication and synthesis; supported by enzymes, transporters and messenger molecules with faster reaction kinetics; both requiring inorganic catalysts to cross the energy barriers to form reaction products, that would ultimately drive the entire system into a ‘steady state’ or ‘homeostasis’^7^. Williams and Rickaby^7^ indeed draw our attention to the weaknesses of the evolutionary theory proposed by Darwin, who had regarded evolution as simply being driven by the ‘survival of the fittest’ rule, as opposed to being defined in the context of the chemical environment in which we can observe changes in the organisms and their environment simultaneously since both are impacted by the other. Indeed, organisms could be viewed as ‘chemotypes’ as opposed to ‘phenotypes’ adapted to the unique bio-geochemical ‘niches’ in which they might exist.

It is hypothesized that the process of translation was pre-dated by the synthesis of polypeptide sequences as quasi-statistical assemblies undergoing initial nucleation and subsequent polymerisation utilizing only a fraction of the modern 20 amino acid code, designated as the ‘early amino acid code’^8,9^. The primordial code was predicted to consist of ten amino acids derived from various sources such as the primitive crust, meteorite bombardment that later resulted in the formation of the ‘late and complete 20 amino acid code’ by the genetically encoded biosynthetic pathways^9^.

Although it is impossible to go back in time to analyse the biogeochemistry of early earth, clues regarding the nature of the earliest functional proteins and their chemical environment can be inferred by measuring the amino acid biases present in proteins encoded by ‘essential genes’ in the ‘minimal protocells’ that could have comprised the LUCA. The rationale of this assumption is that amino acids that arose earlier would be used more frequently within proteins of the LUCA^10^. This might reflect the order of addition of these two groups of amino acids to the genetic code. Although Brooks *et al.*^10^ have attempted to deduce the ‘early amino acid code’ applying ancestral phylogenetic reconstruction techniques in combination with Bayesian modelling techniques to simulate the evolutionary molecular clock, their results are expected to be biased due the unique proteins set used to train their algorithms.

In this paper, the focus is solely on the 30S and 50S ribosomal subunit proteins from two gram negative bacteria- *Thermus thermophilus* and *Escherichia coli*. These two bacteria were selected due to their contrasting biogeochemical niches. Currently, we can speculate that heterogenous and unique LUCA populations could have evolved between −10°C to 150°C in varied habitats across the universe^7^. Observed differences in amino acid frequencies could be then attributed to their unique biogeochemical habitats prevalent on planet earth and on other habitats in our solar system and extended universe.

*T. thermophilus* is a thermophile isolated from hot springs in Japan whose growth temperature is between 47°C to 65-72°C^11^. *E.coli* on the other hand is observed to occur in the open environment as diverse populations including commensal and pathogenic strains on human or animal hosts that can survive between 37°C +− 7°C^12^. Our comparative analysis of amino acid frequencies of *T.thermophilus* and *E.coli* ribosomal proteins with universal UNIPROT^13^ average frequencies, taken as a control set representing the universal amino acid code, reveals remarkable similarity in their respective signatures and key differences. These results indicate that although ribosomal proteins could have originated in LUCA populations in varied habitats, their sequences and amino acid codes have been remarkably conserved during billions of years of evolution possibly due to their critical function in protein synthesis. A temperature sensitive and polarity dependent amino acid code is proposed to explain the origin and evolution of ribosomal proteins. In addition, a nucleation model for ribosome construction involving stepwise accretion of the amino acid code during the evolution of this protein translation machinery complex is described. These results provide critical insights regarding the potential development of the genetic code and the translation machinery. Rules discovered herein could be potentially applied to the systematic study of various molecular complexes in different compartments of eukaryotic cells that arose from a presumably symbiotic relationship of multiple prokaryotes and polymeric assemblies.

## Results

### Amino acid frequencies in 30S and 50S ribosomal proteins provide clues regarding potential sulphur free and anoxic origins of protein translation machinery

Average amino acid frequencies were estimated for all known proteins from UNIPROT spanning ~5631 reference proteomes representing diversity spanning all major taxa such as archaea, bacteria and eukaryotes (Figure 1A and 1B)^13^. This was designated as the normalized control set representing observed frequencies of amino acids in the modern ‘universal amino acid code’. Amino acid frequencies were also calculated for 30S and 50S ribosomal proteins from *T. thermophilus* and *E.coli* and averaged. It was observed that averaged amino acid frequencies corresponding to both 30S and 50S ribosomal proteins showed remarkable similarity to the observed UNIPROT ‘universal average’ except for key amino acid residues such as asparagine, aspartic acid, cysteine, leucine, serine, threonine and tryptophan, that were under-represented; while arginine, glycine, lysine and valine were over-represented. Interestingly, there were species specific differences detected between *T.thermophilus* and *E.coli* with respect to arginine, lysine and proline residues that showed greater abundance in *T.thermophilus* as compared to *E.coli*.

**Figure 1:**
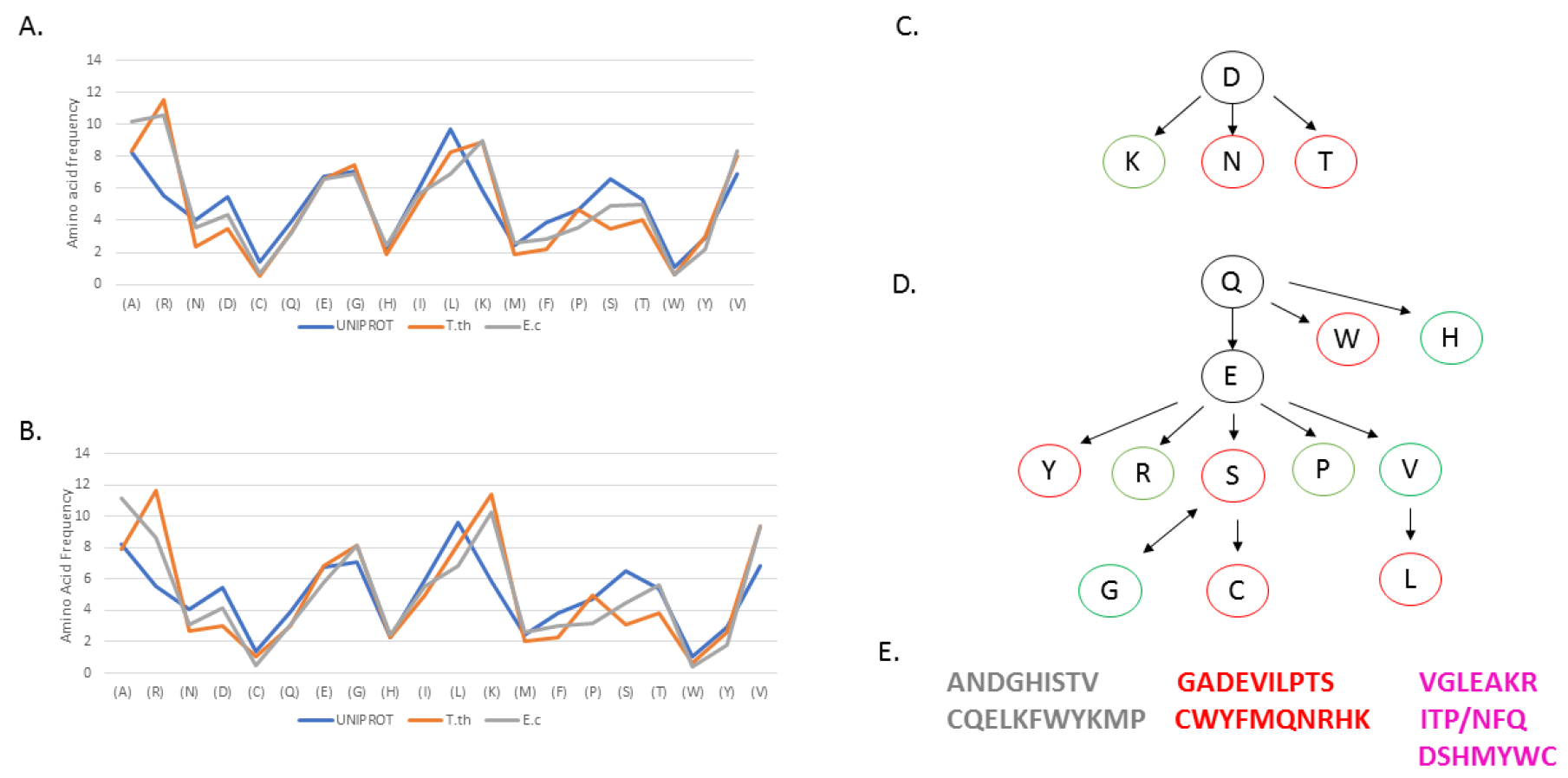
Amino acid frequencies in 30S (A) and 50S (B) ribosomal sub-units of *Thermus thermophilus* and *Escherichia coli*. Average UNIPROT amino acid frequencies were also plotted as a control set. Amino acid bio-synthetic pathways (C) and (D) were constructed from ECOCYC database^17^ where ancient amino acids are shaded red while late amino acids are shaded green. Early (top), middle and late (bottom) amino acid codes as designated by Brooks *et al.*^10^ (grey), Trifonov^8^ (red) and Tripathi (lavender) (E).

Raggi *et al.* have suggested the possibility that primitive RNA-binding protein assemblies could have complexed with negatively charged amino acids such as aspartic acid and glutamic acid associated with metallic cations such as Mg^2+^/Fe^2+^ ^6^. However, stark underrepresentation of negatively charged amino acids such as aspartic acid, which also serves as a pre-cursor to asparagine and lysine residues in the amino acid synthesis pathways (Figures 1C and 1D) and enrichment of positively charged residues such as lysines and arginines indicates that the latter might have played a key evolutionary role in stabilising ribosomal RNAs such as the 23s, 16s and 5s rRNA species that provide the machinery for protein translation (Table 1). These results also hint towards the possibility of aspartic acid independent origin of lysine in the ‘primitive amino acid code’.

**Table 1.**

| T.th/E.c | Function |
| --- | --- |
| S1 | RNA chaperone, unfolds 5'UTR mRNA, autogenous repressor |
| S2 | 16s rRNA binding, S1 interaction |
| S3 | mRNA helicase |
| S4 | 16s rRNA binding, translation, transcription, mRNA helicase |
| S5 | mRNA helicase |
| S6 | 16s rRNA binding, autoregulation of 5'UTR |
| S7 | IF-3 interaction, mRNA binding, rRNA binding, ribosomal structure |
| S8 | translational repression, 16srRNA binding, ribosome assembly |
| S9 | tRNA binding, ribosomal assembly |
| S10 | Transcription anti-termination, tRNA binding |
| S11 | IF-3 interaction, 16srRNA binding, ribosomal assembly |
| S12 | 16s rRNA binding, tRNA binding, splicing |
| S13 | 16s rRNA binding, 50S bridge, tRNA binding |
| S14 | 16s rRNA binding, Zn cofactor (T.th), ribosomal assembly |
| S15 | 16s rRNA binding, 23s rRNA binding |
| S16 | Endonuclease, DNA binding, structural role |
| S17 | 16s rRNA binding, translational efficiency, structural role |
| S18 | 5'UTR binding, 16s rRNA binding, structural role |
| S19 | 23s rRNA binding, tRNA binding |
| S20 | ornithine decarboxylase inhibitor, 16s rRNA binding, structural role |
| S21 | rRNA binding, structural role |

Cysteine has been defined as a sulphur scavenging amino acid that lies downstream of serine biosynthetic pathway. It also plays a key role in forming disulphide bridges in proteins in an oxidizing environment. A systematic deficiency of cysteine residues in ribosomal proteins indicates that these proteins could have originated in anoxic and reducing ecosystems potentially deriving energy for translation from light and U.V. driven electron transfer reactions. An underrepresentation of easily oxidizable amino acids such as methionine, tryptophan, tyrosine and histidine in ribosomal subunit proteins also provide evidence regarding the origin of these complexes in a reducing environment as was prevalent on early earth.

Tryptophan is an amino acid with an aromatic side chain underrepresented in nearly all sampled small and large ribosomal proteins along with cysteine. It’s major biochemical function in higher organisms involves hormonal synthesis along with stabilising membrane proteins. Its exclusion from ribosomal proteins could indicate that the simplest LUCA had minimal membrane proteins and signalling requirements, features that arose later during the course of evolution as living systems evolved in complex chemical environments.

Subtle interspecies differences between *T.thermophilus* and *E.coli* amino acid frequencies are also observed, with arginine, lysine and proline residues being overrepresented in *T.thermophilus* in comparison with *E.coli*. An abundance of proline residues in *T.thermophilus* compared to *E.coli* could also indicate increased structural stability and thus increased evolutionary advantage by this amino acid at higher temperature^14^. Curiously, an absence of glycine residues from L33 in *T.thermophilus* might indicate that this protein could have evolved under high U.V irradiation known to destabilise this amino acid^15^. Species specific differences in amino acid compositions between thermophilic and mesophilic organisms, such as E=K/Q+H ratio >4.5^16^ and enrichment of arginine and lysine residues have been reported previously and are also observed in our data.

In Figures 1A and 1B it was observed that although amino acid frequencies were remarkably conserved for residues such as glutamic acid, glycine, histidine, tryptophan, tyrosine and valine in the 30S sub-units, distinct variations were observed for alanine, arginine, asparagine, aspartic acid, isoleucine, leucine, lysine, phenylalanine, proline, serine and threonine. ~ two-fold increase in frequency of asparagine residues was observed in *E.coli* and UNIPROT compared to *T.thermophilus*. Asparagine is synthesized from aspartic acid by a genetically encoded L-aspartate: L-glutamine amido ligase in a single step reaction that involves addition of amido group from glutamine utilizing energy from ATP. Although it is not currently possible to exactly date its origin in the evolutionary time frame, one can assume that its origin would be intermediate to aspartic acid and cysteine that require multiple steps for their synthesis (Figure 1C and 1D). Interestingly, its abundance in UNIPROT ‘universal average’ is observed to be ~ double of *T. thermophilus*, indicating how a temperature drop could have potentially increased its stability in proteins and thus its abundance in majority of sequenced organisms. Similar changes are observed for other ‘polar’ amino acids such as aspartic acid and serine.

Figure 2 represents the corresponding heatmaps for 30S and 50S amino acid frequencies respectively. Amino acid signatures common to both 30S and 50S ribosomal subunits revealed higher polarity and low hydrophobicity in the over-represented ‘early’ amino acid code VGLEAKR, with an average GRAVY score of −0.357 (Figure 1E) compared to underrepresented ‘late’ amino acid code CWYMHSD with greater hydrophobicity (average GRAVY score= −0.757).

**Figure 2:**
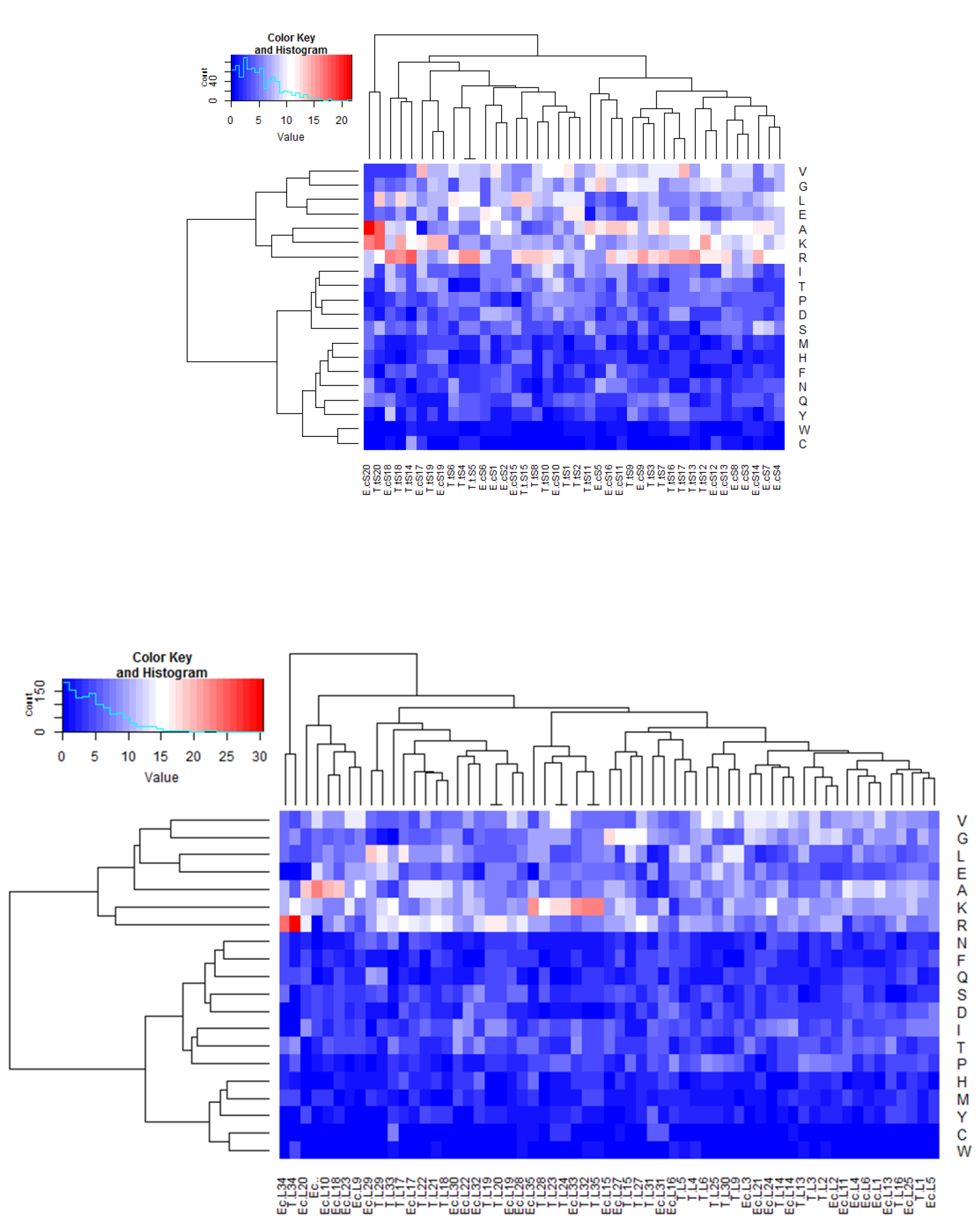
Heatmaps depicting hierarchical clustering of amino acid frequencies were plotted for 30S (top) and 50S (bottom) *T.thermophilus* and *E.coli* ribosomal proteins. Least represented amino acids like C,W,Y,M,H,S,D cluster towards the bottom, while most over-represented amino acids V,G,L,E,A,K,R amino acids lie at the top. NFQ/ITP lie in the middle of the vertically clustered early and late amino acids and are predicted to constitute the ‘middle’ amino acid code. Also see Figure 1E.

### Evaluating different amino acid codes to explain advent of ribosomal proteins

Trifonov provided a multifactorial hypothesis regarding the chronological order of appearance of amino acids using 40 different criteria^8^. G,A,V,D,E,P,S,L,T were designated as ‘early amino acids’ in the chronological order constituting the ‘early amino acid code’. Although origin of the early amino acids such as aspartic acid (Figure 1C) as designated by Trifonov could be explained by a stochastic chemical environment that favoured their formation in tandem with the evolution of nucleic acids such as RNAs, later amino acids are hypothesized to be introduced by the origin of specific genetically encoded biosynthetic pathways, often utilizing early amino acids as precursors. Certain amino acids such as histidine have been found to originate from both inert meteorites as well as from biosynthetic pathways (Figure 1D). Amino acids overrepresented and underrepresented in ribosomal proteins along with their biosynthetic routes derived from ECOCYC database^17^ have been depicted in Figures 1C and 1D.

If ribosomal proteins were truly built using the ancient code as proposed by Trifonov, one would expect a biased enrichment of early amino acids and deficiency of modern or late amino acids (Figure 1E). However, our results demonstrate that underrepresented and overrepresented amino acids in ribosomal subunits are distributed evenly across the early and late amino Trifonov amino acid sets (Figure 1E). Hence, different criteria must be applied to classify the early and late amino acid codes. Brooks *et al.* applied phylogenetic reconstruction and Bayesian modelling^10^ to deduce the early and late amino codes (Figure 1E). However, as with the Trifonov amino acid sets, residues underrepresented and overrepresented in ribosomal proteins are observed to be distributed across both early and late sets. This noise could be attributed to the fact that Brooks *et al.* employ diverse proteins with varied functions to determine conserved residues and build ancestral sequences by phylogenetic reconstruction that could potentially be invalidated by inclusion of paralogs (gene duplications) and horizontal gene transfers during the course of evolution. Hence, the confidence intervals obtained by them to determine the LUCA proteins could potentially generate noisy data with respect to predicting the ‘actual’ amino acid code that specifically led to the origin of the ribosomal proteins.

Intuitively, the problem seems complicated by the fact that although ribosomal proteins show cysteine-tryptophan deficiencies in both the 50S and 30S ribosomal subunits for prokaryotes such as *T.thermophilus* and *E.coli* and eukaryote such as yeast *Saccharomyces cerevisiae* (data not shown), subtle variations in the code are identified for a few selected proteins that could have served as nucleation centres during the origin of the ribosomal complex (Fig 3). For example, L29, L30, L34; L1, L7, L12; S20, S6; and S15 (See Tables 1 and 2) lacking in cysteine, tryptophan, phenylalanine, serine, tyrosine and isoleucine residues in various combinations could have served as nucleation centres binding to 16srRNA, 23srRNA, 5S rRNA, mRNA and tRNAs in the active A and P sites involved peptide bond formation inside the ribosome. This would have been followed by assembly of proteins encoding the ‘modern amino code’ after the evolution of downstream amino acid synthetic pathways, that would have acquired additional regulatory functions during initiation, elongation and termination reactions involved in protein synthesis (Figure 4).

**Table 2.**

| T.th/E.c | Function |
| --- | --- |
| L1 | 23s rRNA binding, tRNA binding, structural role |
| L2 | rRNA binding, peptidyl transferase, 16s rRNA binding, structural role. tRNA binding, Zn binding |
| L3 | 23s rRNA binding, structural role |
| L4 | rRNA binding, peptidyl transferase, 16s rRNA binding, structural role. tRNA binding, Zn binding |
| L5 | 5s rRNA binding, structural role, tRNA binding |
| L6 | 23s rRNA binding, structural role |
| L7/L12 | Binds IF-2, EF-Tu, EF-G, RF3, forms L8, protein homodimerisation, structural role |
| L9 | Binds to 3' end of 23S rRNA |
| L10 | Forms ribosome stalk, negative regulator of translation, forms L8, structural role |
| L11 | 23S rRNA binding, structural role, stringent response, translation termination |
| L13 | Binds 3'end of 23S rRNA, assembly, Zn ion binding, mRNA binding, structural role |
| L14 | Binds 23S rRNA, 16S rRNA, forms bridge between two sub-units, represses translation |
| L15 | Binds 5S rRNA, late assembly |
| L16 | Binds 23S rRNA, located at A site, tRNA binding, assembly, structural role |
| L17 | Ribosomal assembly with L15 |
| L18 | 5S rRNA attachment, increases L5 binding and cooperation with 23s rRNA |
| L19 | Binds close to 5'-end of 23S rRNA |
| L20 | Binds to 5'end of 23S rRNA, important during early stages of 50S assembly |
| L21 | Binds 23S rRNA in the presence of L20 |
| L22 | Binds 23S rRNA with L4, L17, L20, important for assembly. |
| L23 | Binds 23S rRNA , SRP, essential for growth, chaperone docking site, |
| L24 | Assembly protein |
| L25 | Binds 5S rRNA |
| L27 | ribosome binding, rRNA binding, structural role, tRNA binding |
| L28 | rRNA binding, structural component |
| L29 | Binds 23S rRNA, surrounds polypeptide exit tunnel |
| L30 | Structural role, located near GTPase centre |
| L31 | Located near GTPase centre |
| L32 | Structural role |
| L33 | Structural role |
| L34 | Structural role, tRNA binding |
| L35 | Structural constituent |

**Figure 3:**
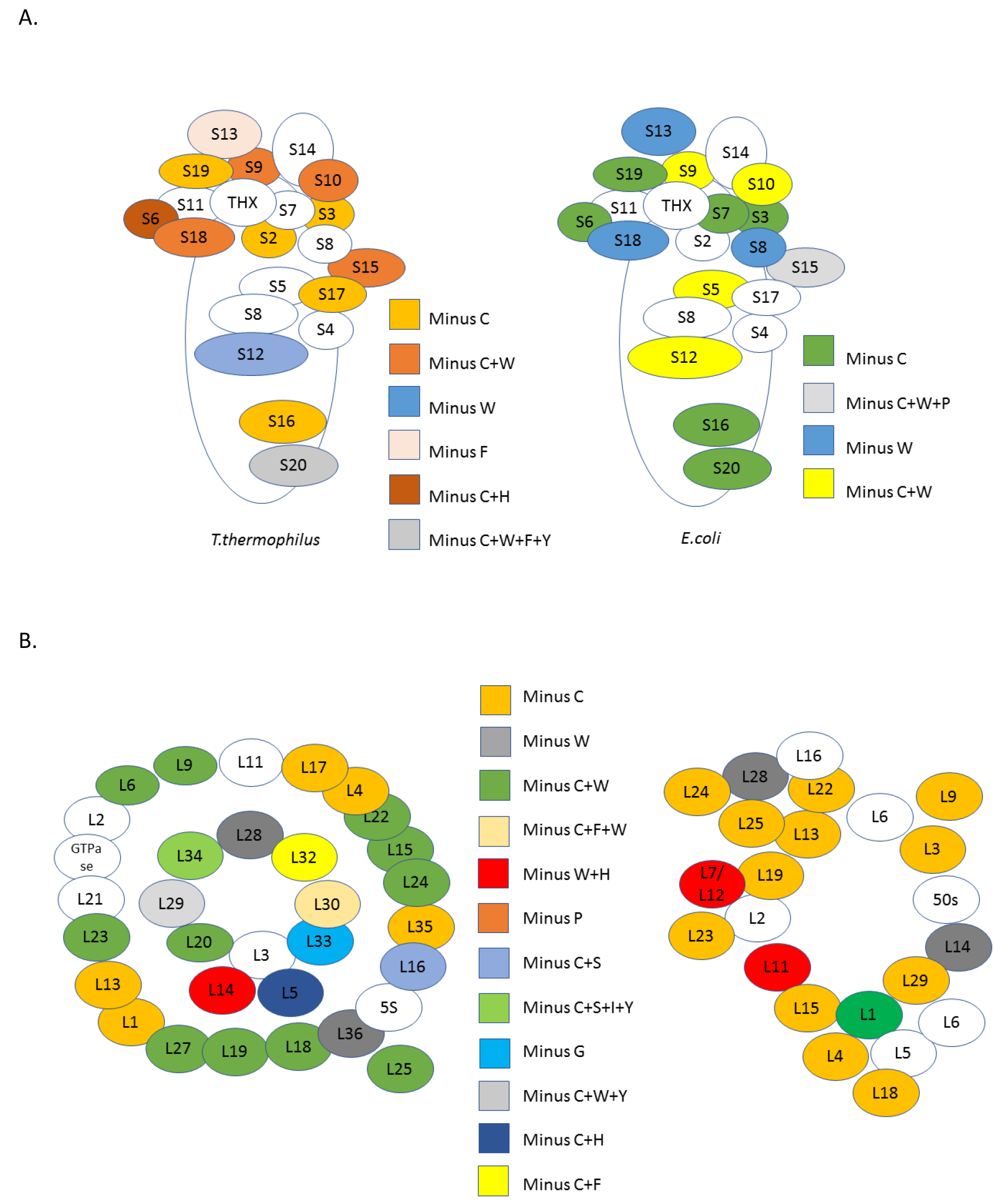
Molecular organisation of 30S (A) and 50S (B) ribosomal sub-units from *T.thermophilus* (left) and *E.coli* (right) were constructed. Proteins with specific amino acid compositions have been colour coded.

**Figure 4:**
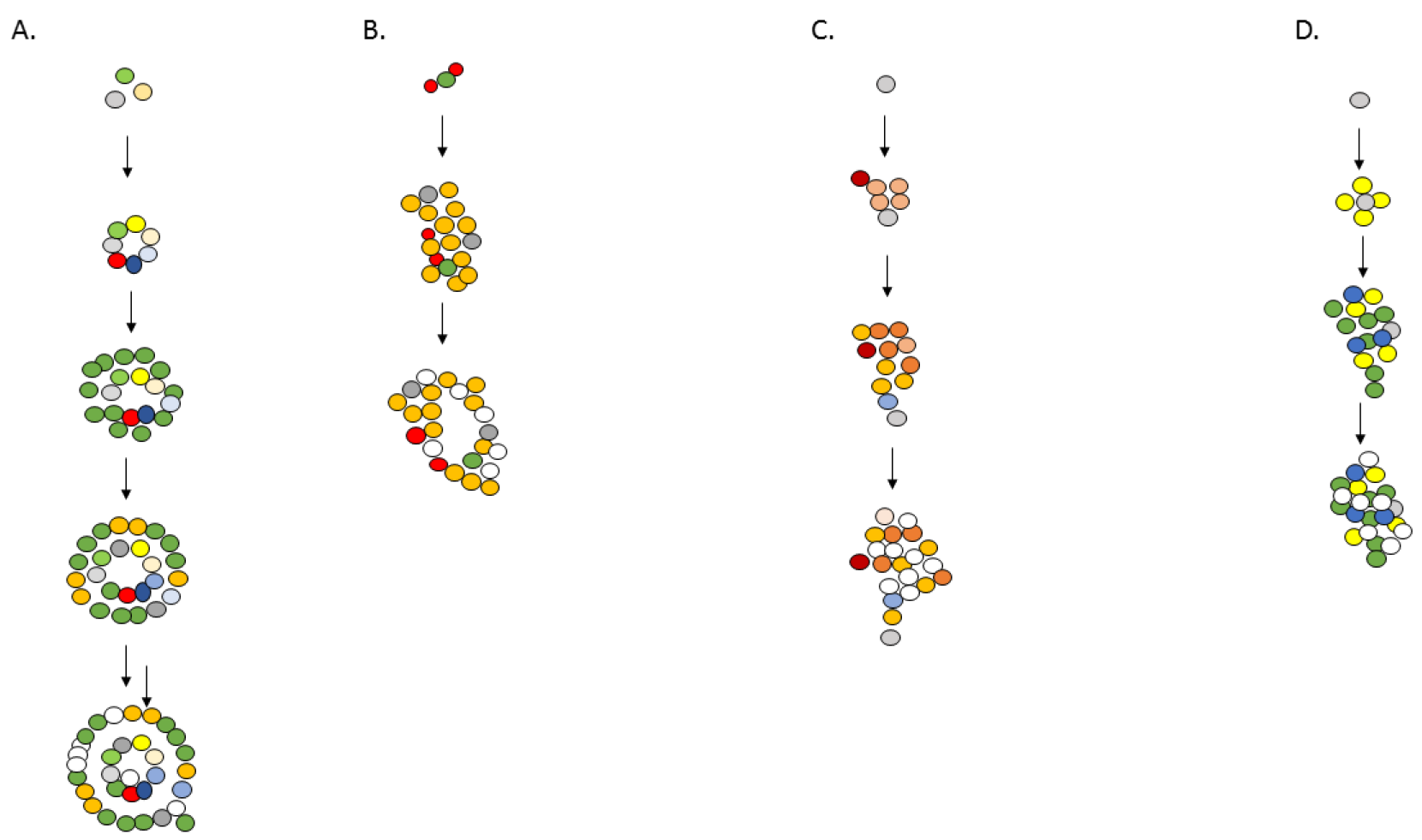
Hypothetically, proteins with maximum amino acid deficiencies would have originated earlier in the evolutionary time-frame. Hence nucleation models for *T.thermophilus* and *E.coli* 50S (A and B) and 30S (C and D) were constructed. Step-wise accretion models that take into account increasing complexity of the ribosomal complexes along with the evolving amino acid code are depicted.

Classification of amino acids into ‘early’ and ‘late’ categories is expected to be marked by some ambiguity due to the scarcity of geochemical data available to us from the earliest periods of planetary formation, when water first condensed on proto-earth, whose environment was highly dynamic, marked by heavy meteorite bombardment and volcanic eruptions. Our understanding regarding when and how amino acids could have potentially arisen presumably in the time spanning the Hadean and the Archean era is currently limited by the contentious nature of fossil records of the earliest microbes and their chemical history, bound to be affected by unknown environmental factors. Therefore, it is assumed that the ‘early’ (VGLEAKR) and ‘late’ (CWYMHSD) ribosomal amino acid signatures deduced in this paper (Figure 1E) are an outcome of the random selection of *T.thermophilus* and *E.coli* species used in our analysis. ITP/NFQ amino acids are ranked in the middle in the vertical rankings deduced in heatmaps corresponding to 30S and 50S amino acid frequencies (Figure 2) and could potentially be classified as the ‘middle’ amino acids in the order of their potential origin in the evolutionary time-frame.

### Step-wise nucleation model for ribosomal construction and accretion of amino acid code

Figures 3 and 4 depict the molecular organisation and step-wise nucleation model of *T.thermophilus* and *E.coli* ribosomal complexes respectively. It is hypothesized that proteins with maximum amino acid deficiencies representing the most ancient amino acid codes would have originated first in the evolutionary time scale in association with 16s, 23s, 5s and tRNA residues. This hypothesis was also supported by analysing the functional data of the 50S and 30S sub-units as identified in literature (Tables 1 and 2). For example, S20 in *T.thermophilus* and S15 in *E.coli* could have initiated nucleation with additional proteins in the 30S sub-units, while L29, L30, L34 in *T.thermophilus* could have initiated a similar accretion in the larger 50S sub-unit. Due to the inability to obtain the complete structural data for *E.coli* proteins only a partial 50S model has been depicted.

## Discussion

This paper provides novel chemical insights into the origin of the ribosomal proteins in *T.thermophilus* and *E.coli* taking into account their unique biogeochemical environments and how their amino acid biases were shaped by temperature, polarity and U.V. irradiation. ‘Early’ and ‘late’ amino acid codes are proposed to have been shaped primarily by the thermostability of polar amino acids such as asparagine, aspartic acid and serine whose frequency is observed to ~ double in mesophilic *E.coli* compared to thermophilic *T.thermophilus*. One can predict a temperature dependency in the evolution and selection of the earliest amino code that could have initially originated at relatively higher temperatures as was prevalent on early earth. A series of complex biogeochemical changes during the co-evolution of earth’s physico-chemical environment are believed to have led to the emergence of the first “anaerobic” life forms^7^. Although it is hypothesized that early environment of earth underwent a transformation from reducing to oxidative where certain chemical gradients might have been formed by natural processes involving energy of the sun, sporadic volcanoes alongside weathering of rocks and minerals by natural geochemical forces, the probability of these chemicals coming together in ‘trapped complex systems’ is not known^7^.

Precursors to first information polymers such as RNA, DNA, proteins and sugars could have potentially initiated from the reaction of simple compounds such as H_2_, CO, NH_3_ and CO_2_ and subsequent cyclisation before 3.5Ga using energy derived from solar radiation, hot reservoir of heat at the Earth’s Centre and chemical catalysts. Other thermal fluctuations (such as day to night variations in temperature, Snowball Earth conditions between 2.4 and 2.1 Ga^7^) could have catalysed slow reaction kinetics driving polymerisation of DNA/RNA/polypeptides. The late advent of amino acids such as cysteine and methionine, as characterised by their low frequencies in protein sets described in this paper, might indicate that primitive anaerobic organisms might have utilised sulphur as H_2_S during photochemical reactions generating energy for the cells under anaerobic conditions. The generation of oxygen around 2.5Ga to about 1.0Ga^7^, potentially enabled the incorporation of sulphur in other organic molecules as S-S and -SH groups, maintaining the redox potential of cells and also allowing novel functioning of transporters, cofactors, coenzymes and metal chelators.

Carl Woese had first proposed the origin of translation systems in a pre-cellular environment due to “direct-templating” between polynucleotides and polypeptides^18^. Nucleobase affinity analysis of mRNAs and both ribosomal and non-ribosomal proteins further suggests ‘direct’ templating of ribosomal proteins occurred before other types of proteins were translated, bridging the gap between the RNA world^19^ and cellular living systems^20^. It is speculated that the earliest amino acids incorporated in 30S and 50S sub-units in *T.thermophilus* were polar due to the origin of these proteins in water along with RNA assemblies. ‘Late’ amino acids are indeed marked by increased hydrophobicity, that potentially arose with the encapsulation of translation systems in membrane enclosed cellular environments. Increased hydrophobicity would have also shaped evolution of the peptide transferase centre (PTC) and the polypeptide exit tunnel in the ribosomal cores, thus enabling the transition of the earliest translation systems from a polar environment into hydrophobic membrane enclosed cellular systems.

Amino acid codes could have indeed evolved distinctly in different habitats to allow unique adaptation and selection of molecular assemblies to specific biogeochemical niches. Understanding the fundamental rules of evolution of the genetic and amino acid code might enable the discovery of potential life forms elsewhere in the universe. The advent of synthetic biology might even allow the unique design of artificial genetic and amino acid codes uniquely suited to extra-terrestrial habitats in the future.

## Materials and Methods

Protein sequences corresponding to 30S and 50S ribosomal sub-units of *T.thermophilus* and *E.coli* were downloaded from NCBI. Amino acid compositions and GRAVY scores were calculated using ProtParam tool on ExPASy^21^ Bioinformatics Resource Portal. Average amino acid composition of UNIPROT entries was taken as a control set for comparison. Graph was plotted using MS Excel and hierarchical clustering of amino acid frequencies was performed using R Bioconductor package^22^.

## Acknowledgements

We would like to thank IIT Delhi for library access and Dr. Ranjana Tripathi for financial support.

## References

1. Glansdorff, N., Xu, Y. & Labedan, B. The Last Universal Common Ancestor: emergence, constitution and genetic legacy of an elusive forerunner. Biol Direct 3, 29 (2008).

2. Kyrpides, N., Overbeek, R. & Ouzounis, C. Universal protein families and the functional content of the last universal common ancestor. J Mol Evol. 1999; 49:413–423. J. Mol. Evol. 49, 413–423 (1999).

3. Miller, S.L. Production of Amino Acids Under Possible Primitive Earth Conditions. Science. 117(3046), 528–9 (1953).

4. Miller, S.L. Production of some organic compounds under possible primitive Earth conditions. J. Am. Chem. Soc. 77, 2351–2361 (1955).

5. Martins, Z. & Sephton, M. Extraterrestrial amino acids. Amino Acids, Peptides and Proteins in Organic Chemistry. 1, (2009).

6. Raggi, L., Bada, J. L. & Lazcano, A. On the lack of evolutionary continuity between prebiotic peptides and extant enzymes. Phys. Chem. Chem. Phys. 18, 20028–20032 (2016).

7. Williams, R. & Rickaby, R. Evolution’s Destiny: Co-evolving Chemistry of the Environment and Life. RSC Publ. (2012).

8. Trifonov, E. N. Consensus temporal order of amino acids and evolution of the triplet code. Gene 261, 139–151 (2000).

9. Higgs, P. G. & Pudritz, R. E. A thermodynamic basis for prebiotic amino acid synthesis and the nature of the first genetic code. Astrobiology 9, 483–490 (2009).

10. Brooks, D., Fresco, J., Lesk, A. & Singh, M. Evolution of Amino Acid Frequencies in Proteins Over Deep Time: Inferred Order of Introduction of Amino Acids into the Genetic Code. Mol Biol Evol 19, 1645–1655 (2018).

11. Oshima, T. & Imahori, K. Description of Thermus thermophilus (Yoshida and Oshima) comb. nov., a Nonsporulating Thermophilic Bacterium from a Japanese Thermal Spa. Int. J. Syst. Bacteriol. 24, 102–112 (1974).

12. van Elsas, J., Semenov, A., Costa, R. & Trevors, J. Survival of Escherichia coli in the environment: fundamental and public health aspects. ISME J. 5, 173–183 (2011).

13. Bateman, A. et al. UniProt: The universal protein knowledgebase. Nucleic Acids Res. 45, D158–D169 (2017).

14. Perálvarez-Marín, A. et al. Influence of Proline on the Thermostability of the Active Site and Membrane Arrangement of Transmembrane Proteins. Biophys. J. 95, 4384–4395 (2008).

15. Ten Kate, I. et al. Amino acid photostability on the Martian surface. Meteorit. Planet. Sci. 40, 1185–1193 (2005).

16. Farias, S. & Bonato, M. Preferred amino acids and thermostability. Genet. Mol. Res. 2, 383–393 (2003).

17. Karp, P. D. et al. The EcoCyc Database. EcoSal Plus 6, 1–19 (2014).

18. Woese, C. The fundamental nature of the genetic code: prebiotic interactions between polynucleotides and polyamino acids or their derivatives. Proc Natl Acad Sci U S A. 59, (1968).

19. Gilbert, W. Origin of life: The RNA world. Nature 618, (1986).

20. Cannon, J., Sherman, R., Wang, V. & Newman, G. Cross-species conservation of complementary amino acid-ribonucleobase interactions and their potential for ribosome-free encoding. Sci. Rep. 5, 18054 (2015).

21. Gasteiger, E. et al. Protein Identification and Analysis Tools on the ExPASy Server. John M. Walk. Proteomics Protoc. Handbook, Humana Press (2005). 571–607 (2005).

22. Ihaka, R. & Gentleman, R. R: A Language for Data Analysis and Graphics. J. Comput. Graph. Stat. 5, 299–314 (1996).

